# *Enterobacter cloacae* complex ST-171 Isolates Expressing KPC-4 Carbapenemase Recovered from Canine Patients in Ohio, USA

**DOI:** 10.1101/337428

**Authors:** Joshua B. Daniels, Liang Chen, Susan V. Grooters, Dixie F. Mollenkopf, Dimitria A. Mathys, Barry N. Kreiswirth, Thomas E. Wittum

## Abstract

Carbapenem resistant *Enterobacteriaceae* (CRE) have emerged as a critical public health threat. Organisms expressing the *Klebsiella pneumoniae* carbapenemase (KPC) were first recognized in the US in the late 1990s and continue to be the predominant CRE genotype reported in clinical isolates. Strains harboring *bla*_KPC_ alleles have been observed in multiple species of *Enterobacteriaceae*, including the *Enterobacter cloacae* complex. A major *E. cloacae* clone, *Enterobacter xiangfangensis* ST171, has emerged as an important cause of hospital associated infections (HAI) and has been shown to carry different alleles of KPC in the context of Tn4401, residing on plasmids of multiple incompatibility groups. While CRE are commonly isolated from infected humans, their recovery from animals has been rare, particularly from companion animals. In the US, only six CRE have been reported from companion animals, and one from livestock, none of which were *bla*_KPC_. This report describes two *E. xiangfangensis* sequence type ST171 isolates each with a large IncHI2 plasmid bearing *bla*_KPC-4_ recovered from dogs with infections at the Ohio State University Veterinary Medical Center. Our phylogenetic comparison of these canine isolates with available sequences from clinical human isolates of KPC-4 identified in ST171 suggest an epidemiologically significant clonal strain.

## INTRODUCTION

Carbapenem class antimicrobial agents are critically important to human health and are considered drugs of last-resort. Carbapenem resistant *Enterobacteriaceae* (CRE) are a growing threat to global public health. In 2013, the CDC classified CRE as an “urgent” public health threat, and estimated that there were approximately 9300 CRE infections, and 610 related deaths in the US (1).

Carbapenem resistance among clinically-relevant *Enterobacteriaceae* is often conferred by plasmid-borne β-lactamase encoding genes that may be categorized in three of the four Ambler β-lactamase classes. KPC and OXA carbapenemases respectively belong to classes A and D and function as serine proteases whose activity is inhibited by agents such as clavulanate and sulbactam. The NDM, VIM and IMP enzymes are class B metallo-β-lactamases which are inhibited by the presence of EDTA, but not clavulanate and sulbactam (2). In the US, the *K. pneumoniae* carbapenemase (KPC) emerged in the late 1990s and have become the predominant carbapenemase encoded by clinical CRE isolates primarily due to clonal dissemination of strains of *K. pneumoniae, Enterobacter spp*., and *Escherichia coli* (3, 4). While healthcare associated infections (HAI) comprise the majority of cases involving CRE overall, community associated CRE infections (CAI) comprise up to 10.8% of infections in the US. Active surveillance performed by hospitals as part of infection control programs indicate that asymptomatic carriage may be up to 41%; however, the rate of carriage in healthy individuals remains unknown (5).

Antimicrobial-resistant bacteria and resistance genes are shared among human beings and animals through either direct or foodborne zoonotic transmission. As a result, veterinary use of antimicrobial drugs in both food-producing and companion animal species has the potential to impact antibiotic resistance of human pathogens (6). While carbapenem antimicrobial drugs are illegal to use in food-producing animals in the US, there is relevant selective pressure favoring CRE in production animal niches resulting from the use of the third-generation cephalosporin, ceftiofur, and other agents that may co-select for carbapenemase encoding genes due to co-localization with other resistance-encoding genes on plasmids (7, 8). To date, there have been two reports of multiple CRE genera expressing *bla*_IMp-64_ located on an IncQ1 plasmid that were recovered from the same US swine farm (9, 10).

In US companion animals (e.g. dogs and cats), it is permissible for a veterinarian to prescribe carbapenem-class drugs off-label, as long as there is a valid veterinarian-client-patient relationship under the Animal Medicinal Drug Use Clarification Act of 1994 (AMDUCA; (11)). While an assessment of the frequency of carbapenem usage in companion animals has not been reported, pharmacokinetic studies have been performed and companion animal dosage recommendations for imipenem and meropenem are available in veterinary medical textbooks (12–14). Moreover, ESBL and AmpC β-lactamase mediated resistance among *Enterobacteriaceae* isolated from dogs and cats is commonly reported, indicating a potential clinical justification for carbapenem administration (15–20).

This potential for the veterinary application of carbapenem drugs suggests that the role of companion animals as reservoirs for the zoonotic transmission of CRE should be considered, particularly for CAIs (21). There has been only one report of CRE isolated from companion animals in the US, six *E. coli* expressing *bla*_NDM-1_ recovered from a variety of infections in cats and dogs between May 2008-May 2009 (22). Subsequent analysis by pulsed-field gel electrophoresis (PFGE) indicated that the isolates represented diverse *E. coli* strains, which is consistent with the dissemination of diverse CRE expressing *bla*_NDM_ in human patients.

CRE can be challenging to diagnose in clinical laboratories because they frequently exhibit clinical susceptibility to carbapenem drugs upon routine antimicrobial susceptibility testing via broth microdilution and Kirby-Bauer disc testing (23). Several strategies, both molecular and phenotypic, have been used to increase the sensitivity of detecting carbapenemase production by clinical isolates (24, 25). In this report, we describe two clinical isolates of *E. xiangfengensis* and their mobile genetic elements recovered from canine patients in central Ohio, US. and belong to a disseminated clone that has been responsible for major clusters of human CRE infections in the Northeastern and Upper Midwestern United States (26).

## RESULTS

### Antimicrobial Susceptibility Testing and Detection of KPC

Both isolates, identified by MALDI-TOF as *E. cloacae*, had the same susceptibility pattern (agent, MIC (μg/ml)): Amikacin, ≤ 4; Amoxicillin/Clavulanic acid, >8; Ampicllin, >8; Cefazolin, >32; Cefovecin, >32; Cefpodoxime, >8; Ceftazidime, >16; Chloramphenicol, 8; Doxycycline >8; Enrofloxacin, >4; Gentamicin, 0.5; Imipenem, ≤ 1; Marbofloxacin, >4; Piperacillin/Tazobactam, 16; Pradofloxacin, >2; Tetracycline, >16; and Trimethoprim/Sulfamethoxazole, >4. Notably, they were phenotypically susceptible to imipenem. Both isolates were observed to produce a carbapenemase by a CarbaNP test, and conventional PCR indicated that both isolates harbored *bla*_KPC_.

### Whole Genome Sequence and Annotation of Plasmid Containing KPC-4

Subsequent Kmer analysis of whole genome sequence identified the isolates as *E. xiangfangensis*, which is included in the *E. cloacae* complex. Both isolates were identified as ST171 on MLST, and each contained two plasmids belonging to IncF1B and IncHI2. The IncHI2 plasmids (Figure 1) were 351,806 (pOSUVMCKPC4-1) and 354,256 bps (pOSUVMCKPC4-2), respectively, and were each identified as ST1 on pMLST. The plasmids both contained *bla*_KPC-4_ in the context of Tn4401b (Figure 2), consistent with other reports of *Enterobacteriaceae* expressing *bla*_KPC-4_. Additional resistance genes in the two genomes were: *strAB, aadAl*, *bla*_OXA-129_, *sul1, tet(B)*, and *dfrA21*. Chromosomal analysis also indicated that they both carry resistance genes *bla*_ACT_16_ and *fosA* in the two isolates. In comparison to pOSUVMCKPC4-1, pOSUVMCKPC4-2 has two additional IS3 at nt 62,536 to 63,760 and nt 180,063 to 181,287, respectively. In addition, a 21kb region (nt 94,348-115,575 in pOSUVMCKPC4-2), containing the *strA, strB, aadAl*, *bla*_OXA-129_, *dfrA21, sul1* flanked by two *IS4321*, is inverted in comparison to pOSUVMCKPC4-1. This region was identified as In524 (Figure 3), using the INTEGRALL database (27). The *intl1* is disrupted by the Tn5393 insertion sequence, which encodes genes *strA* and *strB* that confer resistance to aminoglycoside antibiotics. Downstream of *strB* is a truncated integrase gene followed by three antimicrobial resistance gene cassettes, *dfrA21*, *bla*_OXA_129_, and *aadA1*. The 3’ region following the integron harbors genes encoding resistance to quaternary ammonium compounds, *qacEΔ1*, and sulfonamide antibiotics, *sul1*. This region also contains a putative gene for N-Acyltransferase, Acyl-coA, and the insertion sequence IS6100, which is a transposase shown to increase the expression of *strA* and *strB* genes. The IncF1B plasmids in the two isolates were identical in size, 108,403 bp, and contained no identifiable resistance genes.

**FIG 1.**
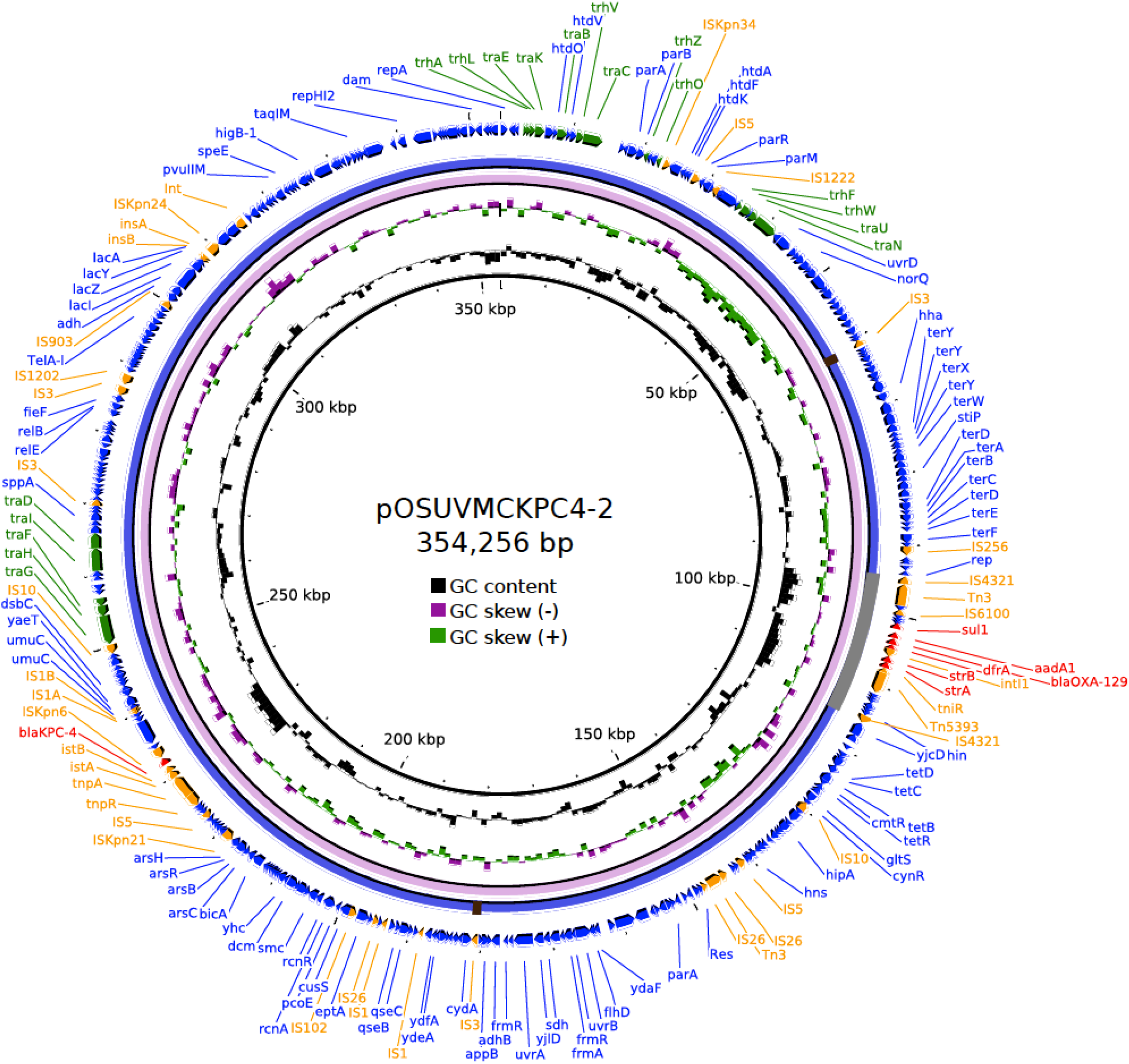
IncHI2 plasmids isolated from *E. xiangfengensis* ST171 sourced from dogs (pOSUVMCKPC4-2 (violet) and pOSUVMCKPC4-1 (blue)). Outer ring denotes location and orientation of genes. Subplasmidic mobile element associated genes are indicated in orange and red and plasmid transfer genes are indicated in green.

**FIG 2.**
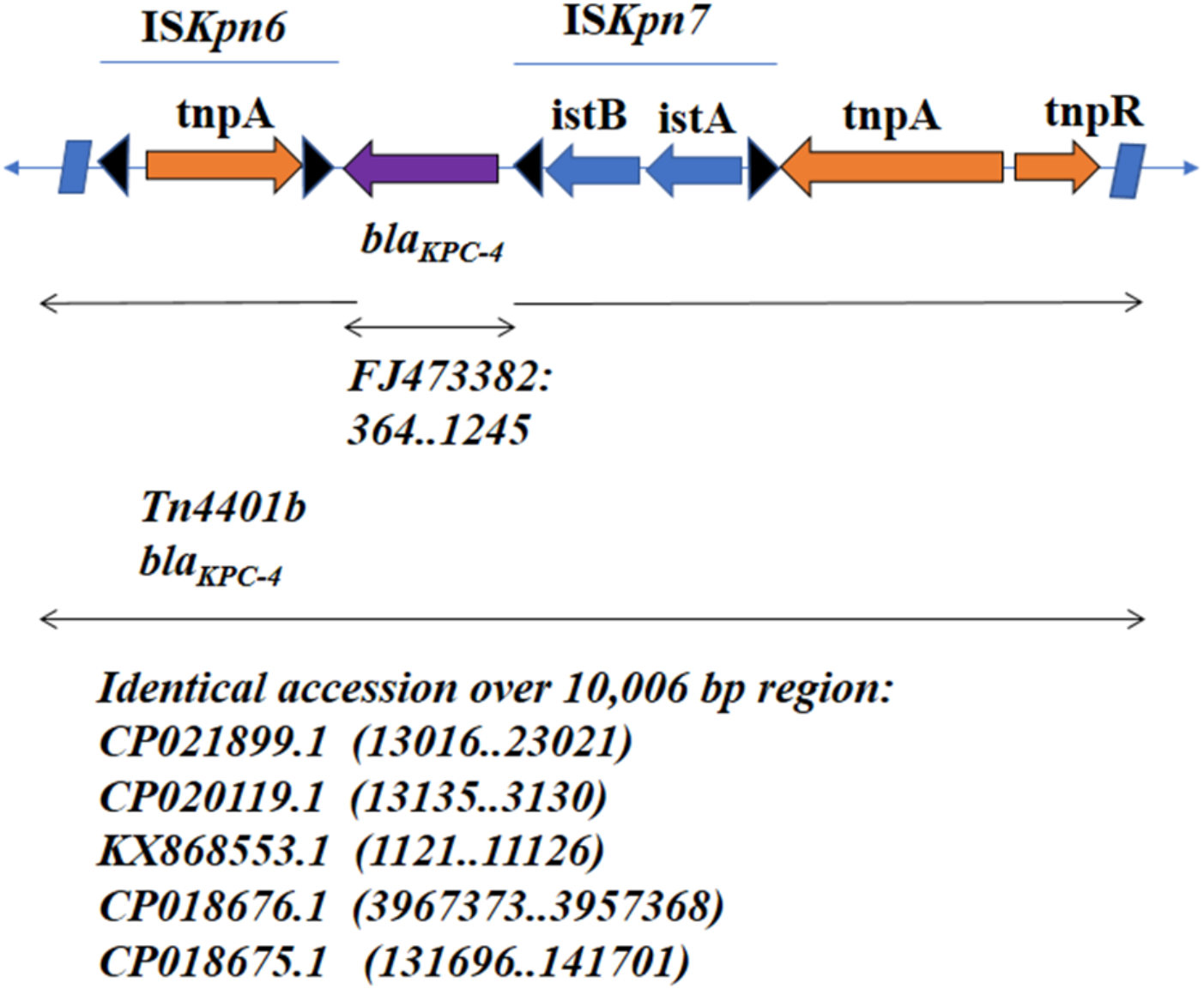
Schematic structure of Tn4401b containing *bla*_KPC-4_ present on large IncH12 plasmids in *Enterobacterxiangfangensis* ST171 recovered from two canine patients of The Ohio State University Veterinary Medical Center in 2016. Genes and their corresponding transcription orientation are indicated by colored horizontal arrows. Corresponding insertion sequences (IS) that encode transposons are indicated above the figure with a narrow blue line. Black triangles indicate inverted repeat right and left sequences associated with the *ISKpn6* and *ISKpn7*. Flanking parallelograms represent the two inverted repeat sequences associated with Tn4401 structures.

**FIG 3.**
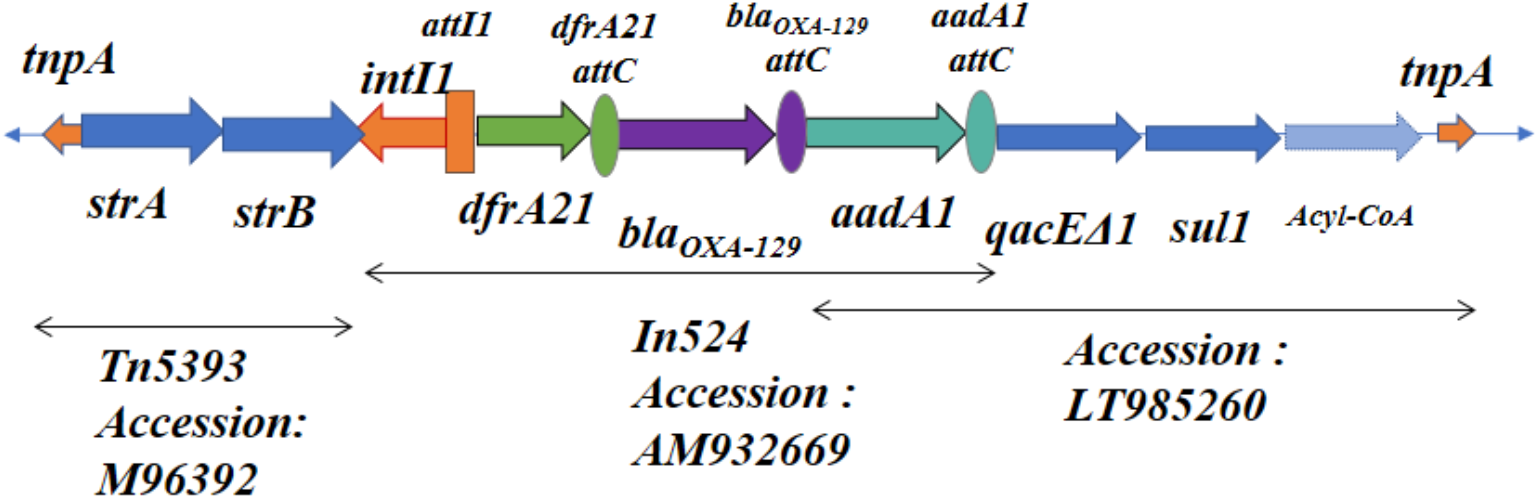
Schematic structure of the In524 class 1 integron and the surrounding region present on large IncH12 plasmids in *Enterobacter xiangfangensis* ST171 recovered from two canine patients of The Ohio State University Veterinary Medical Center in 2016. Genes and their corresponding transcription orientations are indicated by colored horizontal arrows. Homologous alignments to references are indicated with narrow, double sided black arrows with region description and corresponding accession number below. Recombination sites, attC, of gene cassettes are illustrated as ovals with color corresponding to the related gene. Integron gene cassettes are incorporated into the rectangular attI1 site corresponding to the class 1 integron.

Core SNP phylogenetic analysis placed these two *E. xiangfangensis* strains in the previously described ST171 cluster II (Figure 4)(26). The two isolates formed a subclade in cluster II, and differ from each other by 14 core snps. In cluster II, isolates have an average of 75 (10–137) core snp differences compared with each other, while they showed ~530 core snps difference in comparison to isolates from cluster I. Of interest, nearly all isolates from cluster II harbor the *bla*_KPC-4_ on the Tn4401b element. Further analysis of the *bla*_KPC-4_-harboring contigs from other clade II isolates revealed that similar to OSUVMCKPC4-1 and OSUVMCKPC4-2, the majority (except for SMART-264 and BIDMC94) harbor *bla*_KPC-4_ and IncH plasmid backbones.

**FIG 4.**
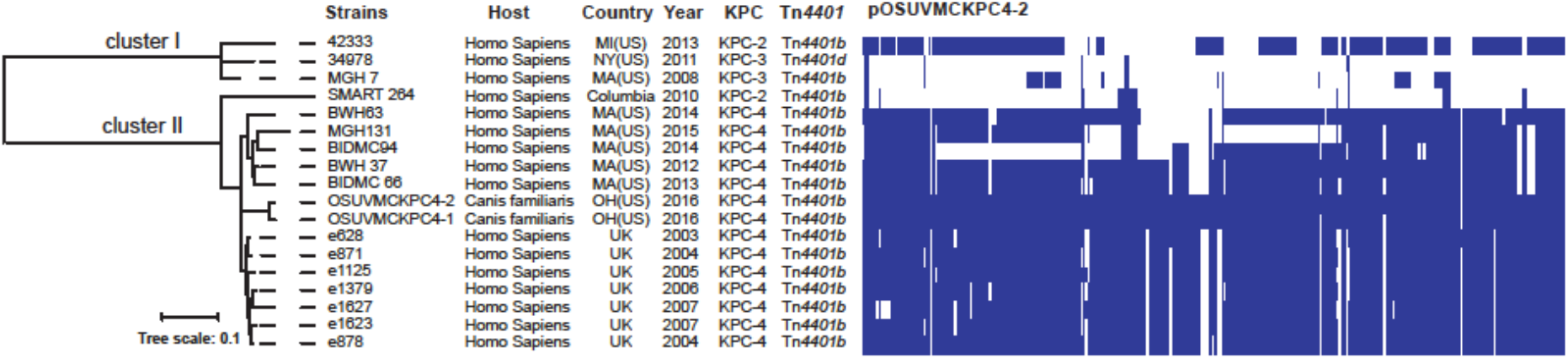
Heatmap of plasmids among KPC-4 bearing *Enterobacter* ST171 with additional KPC-2 and KPC-3 containing ST171 isolates included as references. (Left) Core SNP phylogenetic tree generated by RAxML. (Center) Metadata, including host, isolation location, year, *Tn4401* isoform, and KPC allele. (Right) Plasmid composition is illustrated by showing the BLASTn matches to each *Enterobacter* genome across all of the genes on the pOSUVMCKPC4-2 plasmid.

## DISCUSSION

This first reported isolation of KPC-producing CRE from US companion animals has important implications for both veterinary medicine and public health, because animals may serve as reservoirs of significant opportunistic human pathogens. Notably, these two cases represent an epidemiologically significant clone of *Enterobacter* sp., ST171 that has been described in regional HAI clusters in the US (28, 29), and which may now be expanding into the community. Analysis of contemporaneous human clinical *E. cloacae* isolates from The Ohio State University Medical Center collected between 2011-2016 with an extended-spectrum β-lactamase phenotype (n=8) revealed that two isolates belonged to ST171, and both contained *bla*_KPC-2_ (data not shown), suggesting that these canine strains may have been uniquely community-associated during that time period. However, our SNP analysis comparing the canine isolates to other available ST171 whole genome sequence data in GenBank indicated that they are closely related to historical human clinical isolates from the Northeastern US, the UK, and Colombia. Together, these observations indicate the diversity regarding the dissemination of ST171 strains harboring *bla*_KPC_, including transcontinental movement and circulation in the community over multiple years, with sporadic HAI activity.

The role of direct selection pressure by antimicrobial agents associated with the two infected dogs is unclear. The first dog had a recent history of doxycycline administration for treatment of a chronic pyoderma, which may have selected for the multidrug resistant *Enterobacter*, as the isolate was also tetracycline resistant. However, co-selection for carbapenem resistance by tetracycline antibiotics has not been reported. The second dog had no recent history of antimicrobial drug administration; however, because the organism and the infection was the result of a dog bite from another dog, it is possible that the biting dog had undergone antimicrobial selection pressure and inoculated the open wound with the carbapenem-resistant *Enterobacter*. The dog with the bite wound had not previously been seen at this facility and the organism was isolated from a specimen acquired on the day of presentation, approximately two months after the first dog, thus it is reasonable to conclude that there was not a common nosocomial source. Moreover, medical records indicated that the dogs resided in cities greater than 240 km apart, suggesting that there is significant regional community dissemination of ST171 in Ohio.

Laboratory detection of CRE is a challenge to all clinical diagnostic laboratories, but represents an even a greater challenge to those in the veterinary setting because of the low prevalence in clinical isolates from animals. blaKPC-4 can be particularly difficult to detect with conventional phenotypic (MIC) methods because of its low hydrolytic activity (30), which may be compounded by the weak promoter activity of *TN4401b*, as in these two isolates (31). The isolates described in this study were detected as part of a surveillance program at a tertiary-referral veterinary hospital that is associated with a university. Before the implementation of the surveillance program, only conventional AST would have been performed, and these CRE would not have been detected. Veterinary diagnostic laboratories in the US typically utilize AST performance standards with susceptibility breakpoints that are based on animal pharmacokinetics-pharmacodynamics which are published in the CLSI VET01 document (32) and supplemented with human breakpoints published in the same document when no animal-specific breakpoints are available. While CLSI VET01 does not mention the need for enhanced detection methods for carbapenemase activity, its human counterpart, the CLSI M100S27 document, discusses the need for specific testing (33).

As fewer antimicrobial drugs retain their efficacy in the face of the proliferation of multidrug resistant bacteria, there is a need to take a “One Health” approach that encompasses factors beyond antimicrobial agent usage and infection control in human medicine. The identification of KPC-encoding *E. cloacae* complex organisms in a companion animal species is more evidence that there are biological reservoirs and potentially vectors for transmission to human beings in community circulation. Additional surveillance work and studies of veterinary antimicrobial usage, as well as increasing the capabilities of veterinary diagnostic laboratories will be critical to slowing the dissemination of CRE.

## MATERIALS AND METHODS

### Bacterial Isolates

Both isolates, OSUVMCKPC4-1 and OSUVMCKPC4-2, were originally sourced from clinical veterinary patients (dogs) that were treated at The Ohio State Veterinary Medical Center. The first dog, seen in July 2016 for acute bacterial cystitis (UTI), was a 13-year old female, ovariohysterectomized (spayed) Shetland Sheepdog that had a two-month history of chronic kidney disease and a two-year history of pyoderma. The second dog, seen in September 2016, was a 6-year old male, castrated, mixed-breed dog that presented for an infected bite wound that resulted from a fight with another dog. The first dog was an established patient of the hospital for ongoing management of her chronic health problems and had received doxycycline during the previous year; however, the second dog had never been seen at the facility prior to presenting with the bite wound (which was sampled for culture on the day of presentation), and had no recent history of antimicrobial administration.

### Identification and Antimicrobial Susceptibility Testing

After routine aerobic cultivation from clinical specimens, isolates were identified via MALDI-TOF (Biotyper, Bruker Daltonics, Billerica, MA) and antimicrobial susceptibilities were determined via broth microdilution in accordance with CLSI VET01A4 (COMPGN1F plate, Trek Sensititre, Thermo-Fisher). Carbapenemase production was assessed using a CarbaNP test (34). Isolates were screened for the presence of the transmissible carbapenemase gene, *bla*_KPC_, by conventional PCR using previously reported primers (35, 36).

### Whole Genome Sequencing

Both isolates were first sequenced using Illumina MiSeq and subsequently by the PacBio RS II system at the University of Maryland Institute for Genome Sciences (OSUVMCKPC4-1) and at the University of Delaware Sequencing and Genotyping Center (OSUVMCKPC4-2) using two SMRT cells per isolate, with genomes assembled using Canu v1.4 (37). Isolate DNA was prepared for PacBio sequencing using a commercial DNA extraction kit (Qiagen Blood & Cell Culture DNA Maxi Kit (Qiagen, Germantown, MD). Initially, sequences were submitted to the Center for Genomic Epidemiology website (https://cge.cbs.dtu.dk) for Multi Locus Sequence Typing (MLST), plasmid identification and resistance gene detection using PlasmidFinder and ResFinder (38–40). Additional antimicrobial resistance databases were used to identify genotypes (41, 42). Identification of insertion sequences that encode transposases were identified by IS Finder, as well as annotation of the corresponding IRL, IRR (43). The curated integron database, INTEGRALL (27), identified the class 1 integron. Annotation of plasmid regions of interest were first examined for functional genes with NCBI’s Conserved Domain Database search (44). Regions that contained antimicrobial resistance genes were then used as query sequences to search the NCBI database with BLAST for similar regions (45). Sequences are available in GenBank through NCBI (accession numbers CP024908 and CP029246) and were additionally annotated using the NCBI automated annotation pipeline.

An additional 16 KPC-producing ST171 genomes from the GenBank Whole Genome Shotgun (WGS) database were downloaded and compared OSUVMCKPC4-1 and OSUVMCKPC4-2. A core singlenucleotide polymorphism (SNP) (defined as SNPs shared across all genomes) analysis was conducted using kSNP3.0 (46), and a core SNP maximum-likelihood tree was produced by RAxML 8.2.4 (47) using the GTRGAMMA model and 100 bootstrap replicates. Furthermore, BLASTn comparisons between each isolate’s de novo assembly and the reference pOSUVMCKPC4-2 plasmid were conducted using the method described previously (26), and the phylogenetic tree was annotated using iTOL (48).

